# Ecological divergence of sibling allopolyploid marsh orchids is associated with species specific plasticity and distinct fungal communities

**DOI:** 10.1101/2024.04.24.590886

**Authors:** Katie Emelianova, Anna-Sophie Hawranek, Mimmi C. Eriksson, Thomas M. Wolfe, Ovidiu Paun

## Abstract

Phenotypic plasticity, the dynamic adjustment of traits to environmental variations, is crucial for enabling species to exploit broader niches and withstand suboptimal conditions. This adaptability is particularly relevant for newly formed allopolyploids, which possess redundant gene copies and must become established in diverse environments distinct from their parents and other relatives. By evaluating gene expression and root mycobiome among two ecologically divergent sibling allopolyploid marsh orchids (*Dactylorhiza majalis* and *D. traunsteineri*) in reciprocal transplants at localities where both species are native, we aimed to understand the drivers of species persistence in the face of interspecific gene flow. Despite consistent abiotic differences characterising the alternative environments at each locality, the majority of gene expression differences between the allopolyploids appears to be plastic. Ecologically relevant processes, such as photosynthesis and transmembrane transport, include some genes that are differentially expressed between the two orchids regardless of the environment, while others change their activity plastically in one species or the other. This suggests that although plasticity helps define the specific ecological range of each sibling allopolyploid, it also mediates gene flow between them, thereby preventing differentiation. Extending our investigations to the root mycobiome, we uncover more diverse fungal communities for either species when grown in the environment with nutrient-poor soils, indicating that both abiotic and biotic factors drive the distribution of sibling marsh orchids. Altogether, our results indicate that plasticity can simultaneously promote diversification and homogenization of lineages, influencing the establishment and persistence of recurrently formed allopolyploid species.

**Significance statement:** This study highlights the role of phenotypic plasticity in the persistence and distribution of sibling allopolyploid marsh orchids (*Dactylorhiza majalis* s.l.). By examining gene expression and the diversity of root mycobiome across reciprocal transplantations in native environments, we uncover high plasticity that facilitates adaptation and gene flow, thereby promoting both diversification and homogenization. These findings underscore the importance of plasticity in the establishment and long-term survival of allopolyploid species across the landscape.

## Introduction

Polyploidy, or the duplication of the whole genome, introduces substantial genetic redundancy, which has been recognised as a diversifying force that may enhance the capacity to express different gene copies under varying environmental conditions (de Jong & Adams, 2023). This elevated genetic redundancy is expected to increase phenotypic plasticity (Jackson & Chen, 2010; Mattingly & Hovick, 2023), the ability of an organism to adjust its phenotype to environmental variation and thus track a dynamically changing trait optimum beyond the limits of a fixed genotype (Schlichting & Pigliucci, 1998). However, increased phenotypic plasticity is often regarded as inherently costly as soon as the population has reached an adaptive optimum (DeWitt et al., 1998), in particular in naturally or anthropogenically challenging environments - habitats frequently occupied by polyploids and hybrids, which thereby avoid direct competition with diploid and other relatives. Moreover, some authors have argued that plasticity might slow down the evolutionary process, by reducing the efficiency of natural selection (Charlesworth et al., 1982) or even by resulting in maladaptation (Arnold et al., 2019; Van Kleunen & Fischer, 2005). In stark contrast, some studies have documented adaptive benefits of plasticity, for example related to responses to abiotic or biotic stress (Auld & Relyea, 2011; Nicotra et al., 2015; Solé-Medina et al., 2022), whereas other authors have involved plasticity in initial stages of phenotypic evolution as a key mechanism fostering local adaptation (Mallard et al., 2020; Szukala et al., 2023).

Polyploid species typically originate as small founding populations that face additional challenges associated with minority cytotype disadvantage when sympatric with a majority of diploid individuals (Husband, 2000; Levin, 2002). Polyploids often form recurrently, increasing the genetic diversity of the new lineages (Soltis & Soltis, 1999). If they form polytopically or at different evolutionary times, they may encounter contrasting environments and therefore selective pressures, which may lead to formation of distinct sibling polyploid species (Paun et al., 2007). Phenotypic plasticity can allow polyploids to overcome an initially depauperate gene pool by extending their habitable niche beyond their fixed genotypic capacity (Castillo et al., 2018), at the same time potentially enabling colonisation of environments that are free from homo- or heteroploid relatives (Hahn et al., 2012). However, increased plasticity extends ecological niches, and hence may also facilitate gene flow between establishing polyploids and with their relatives, alleviating in the long term some of the founder effects of low genetic diversity. Studies addressing the role of plasticity in polyploid evolution often focus on the immediate implications of polyploid plasticity, such as whether plasticity is heightened in polyploids compared with diploids (Drunen & Johnson, 2022). However, the role of plasticity in shaping the distribution of young sibling polyploid species remains poorly understood, despite its likely significant implications for how whole genome duplication contributes to biodiversity (Mattingly & Hovick, 2023).

To understand the interplay between plasticity, gene flow and ecological differentiation in polyploids, we focused on the marsh orchids *Dactylorhiza majalis* (Rchb.) P.F.Hunt & Summerh. and *D. traunsteineri* (Saut. ex Rchb.) Soó (Fig. 1a). These are sibling allopolyploids (2*n* = 80) resulting from independent and unidirectional allopolyploidisation events between diploids *D. incarnata* (L.) Soó (the paternal parent in either case) and *D. fuchsii* (Druce) Soó (the maternal parent) (Brandrud et al., 2020; Pillon et al., 2007). Typically, *D. majalis* includes larger plants, with more flowers per inflorescence and broader leaves than *D. traunsteineri* (Fig. 1a). The two orchids have generalist pollination syndromes, corresponding to a food-deception strategy (i.e., no reward is offered to pollinators) (Hansen and Olesen, 1999; Ostrowiecka et al., 2019). It is generally accepted that *D. traunsteineri* is the younger of the two sibling allopolyploids, and estimates of formation place both origins at different times in the recent half of the Quaternary (Brandrud et al., 2020; Hawranek, 2021; Nordström & Hedrén, 2009). Both sibling allopolyploids have a wide European distribution, with *D. majalis* having a more continuous distribution towards southern areas from the Pyrenees to southern Scandinavia, while *D. traunsteineri* occupies disjunct northerly locations in Scandinavia and the British Isles, with their distributions overlapping in the Alps. However, the delimitation of these allopolyploids is debated with different authors recognizing them as subspecies of *D. majalis* s.l. (e.g., Nordström & Hedrén, 2009), as distinct species (e.g., Brandrud et al., 2020; Pillon et al., 2007), or referring to *D. traunsteneri* as an aggregate of several species (e.g., Bateman, 2011). Gene flow between the two allopolyploids has been documented in areas of sympatry, with a tendency for more intense gene flow from *D. majalis* to *D. traunsteineri* (Balao et al., 2016; Brandrud et al., 2020). However, the sibling allopolyploids maintain distinct ecologies differentiated by soil chemistry, including nutrient content and water availability (Fig. 1; Paun et al., 2011; Wolfe et al., 2023), and both species are found in their respective ecologies even at local scales in regions of sympatry. Despite the documented gene flow, previous studies uncovered ecologically-relevant transcriptomic differences between the allopolyploids (Wolfe et al. 2023; Eriksson et al., 2024), accompanied by a stable epigenetic divergence correlated with eco-environmental variables (Paun et al., 2010).

**Figure 1.**
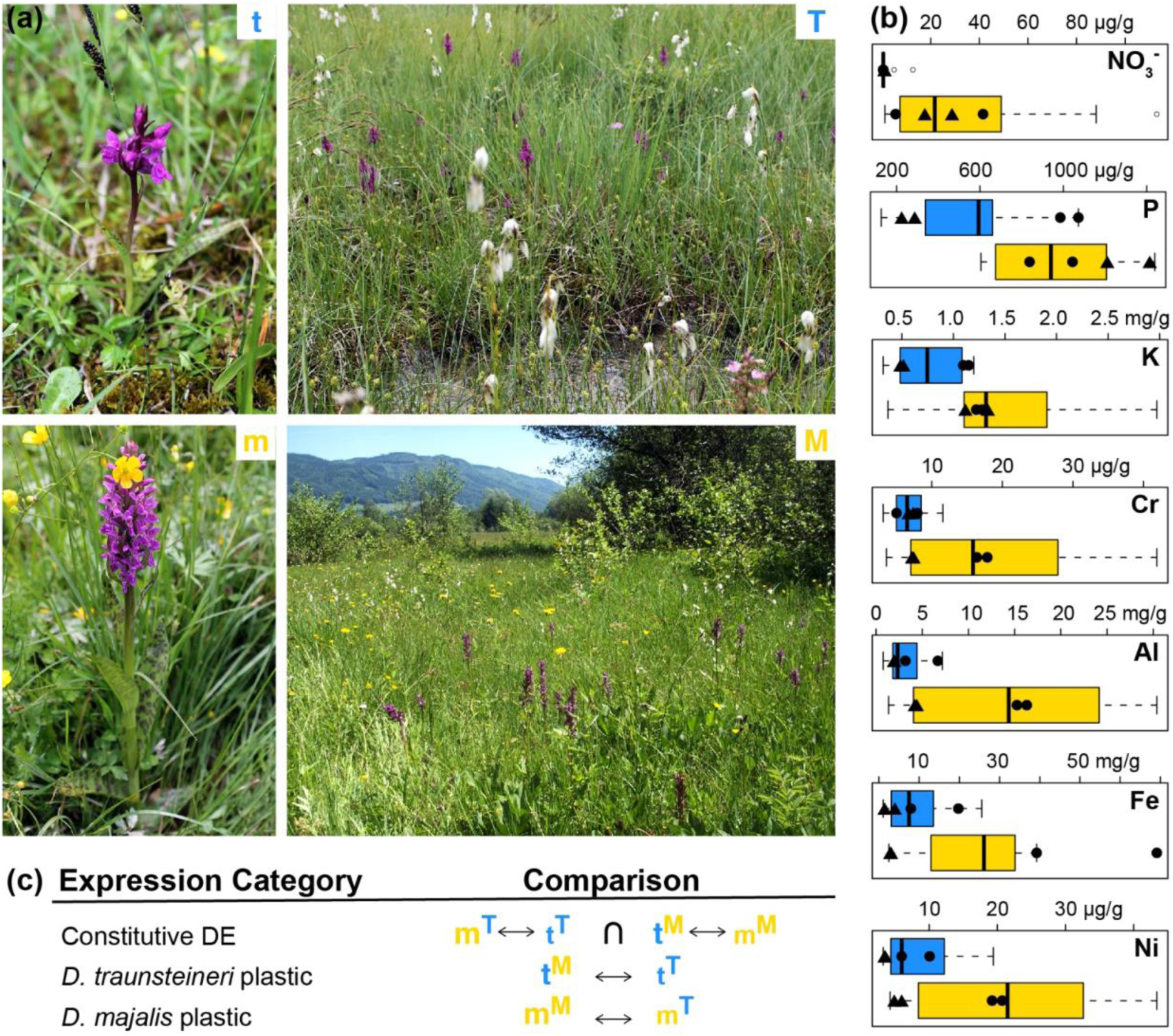
The study system and experiment design. (a) The study species: t, *Dactylorhiza traunsteineri* and m, *D. majalis* growing at one of the transplantation sites at St. Ulrich am Pillersee, Austria. T and M show impressions of the two environments at the same locality. (b) Divergent preference for selected soil characteristics between *D. traunsteineri* (blue) and *D. majalis* (yellow) across multiple European populations (data from Wolfe et al., 2023). With filled circles two measurements per species are indicated at the experimental site at Kitzbuhel, and with triangles at the site at St. Ulrich am Pillersee, both Austria. (c) Scheme of main comparisons made in differential expression (DE) testing. As above, lower case letters indicate sample species, superscript capital letters indicate the environment in which the sample was experimentally grown. ↔ indicates test of differential expression between sample groups, ∩ indicates intersection of genes found to be differentially expressed between sample groups.

We investigated in the present study the rates of expression of genes in leaves and roots of *D. majalis* and *D. traunsteineri* at two localities where the sibling allopolyploids naturally coexist in sympatry and experience varying levels of gene flow (Balao et al., 2016; Brandrud et al., 2020). Using a reciprocal transplantation design, we grew individuals from each species in their own environment and in that of the other species to understand how much of their gene expression is genetically encoded versus how much is underpinned by plasticity. This approach enabled us to identify genes with differences in expression rates between the two species independent of the environment, likely representing genes which have become canalised (i.e., contribute to adaptation to the environment of each respective species and are no longer plastic) and may avoid the homogenising effects of gene flow in sympatric populations. Comparing the proportion and function of plastic and constitutive genes in admixed populations enabled us to understand the factors which define the ecological ranges of *D. majalis* and *D. traunsteineri*, and the proportion of the gene space (Dodsworth et al., 2015) involved.

We also investigated the potential for fungal communities to reinforce species boundaries, utilising the reciprocal transplantations to test the consistency of these profiles across independent localities. Mycorrhizal fungi provide vital support, especially during early stages of the developmental cycle of orchids, and can ameliorate effects of nutrient deficiencies in host plants (Bhantana et al., 2021; Saboor et al., 2021). However, the broader role of fungi in determining host plant success and, in general, species boundaries remains poorly understood. Leveraging the capture of contaminants during RNA-seq experiments, we performed fungal taxonomic profiling of root RNA-seq data from native and transplanted samples to test whether the environments of *D. traunsteineri* and *D. majalis* differed in fungal community profiles. Differences in soil fungal communities may represent an important ecologically relevant distinction between the respective habitats (see Wolfe et al., 2023 for detailed abiotic characterizations), suggesting that mycobiomes could play a role in the delimitation of species boundaries.

This work uses *in situ* measurements of gene expression, functional response to different environments, and fungal community profiling to investigate the consequences of transgressing ecological optima between two sibling allopolyploid species. Our final aim is to provide a deeper understanding of how resilience to environmental heterogeneity interplays with ecological adaptation to maintain species boundaries between sibling allopolyploid marsh orchids in the face of gene flow.

## Results

Two localities where the two allopolyploids exist within a few hundred metres from each other have been selected in Tyrol, Austria for transplantation experiments: Kitzbühel and St. Ulrich am Pillersee. For each locality and species, five individuals were transplanted in spring 2016 and grown in the environment of the other species, and five individuals were transplanted and grown in their own environment to control for transplantation effects. After two growing seasons, leaf and root tissues have been sampled to produce and sequence total RNA-seq libraries (see Methods).

### Global effects of transplantation

Leaf tissue libraries consisted on average of 65 ± 30 million pairs of reads, whereas root libraries consisted of 107 ± 18 million. Of these, 43.9% and, respectively, 44.2% mapped uniquely to the available *D. incarnata* v.1.0 reference genome (Wolfe et al., 2023) (see Supplementary Table S1 for further mapping details). The genomes of both *D. majalis* and *D. traunsteineri* have a high content of repetitive elements (estimated at ca 73%; Eriksson et al., 2022) which likely reflects in a high proportion of LTR reads in the ribosomal depleted total RNA data and may account for the decreased rate in uniquely mapping reads.

We evaluated the genetic structure of the randomly selected accessions using principal components analyses (PCA) using pcANGSD v0.99 (Meisner & Albrechtsen 2018). After excluding low confidence variants, those associated with repetitive regions, and keeping only putatively unlinked loci, we retained 315,846 polymorphic positions for this analysis. The first axis of the PCA explaining 4.2% of the variation (Supplementary Fig. 1) separates *D. traunsteineri* accessions from St. Ulrich from the rest, whereas the second axis (representing 1.4% of the variation) separates *D. traunsteineri* accessions from Kitzbühel from the rest. In line with previous results (Balao et al. 2016, Brandrud et al. 2020) *D. majalis* accessions from either locality form a more cohesive genetic cluster, whereas *D. traunsteineri* individuals separate very clearly by locality. Finally, whereas *D. traunsteineri* shows no genetic separation by environment, the samples of *D. majalis* show a minor tendency of separation by environment on the second PCA axis. As the individuals for the experiment were selected by chance, this slight separation may track back to a somewhat different expression rate of this species between the two environments, resulting in slightly different genotype likelihoods

We further used expression-based PCAs to visualise the global transcriptional profiles of the accessions, separated by tissue type and locality (Fig. 2 A and C). The expression profiles tended to group samples by species and by environment, though the trend was weaker in leaf tissue, root tissue showing a stronger separation of species regardless of environment. Considerable variation was present in the expression profiles of conspecific accessions, even among those sampled in the same environment. *Dactylorhiza majalis* accessions were more variable than *D. traunsteineri*, and transplantation separated *D. traunsteineri* individuals to a greater extent than *D. majalis*, especially in leaf tissue.

**Figure 2.**
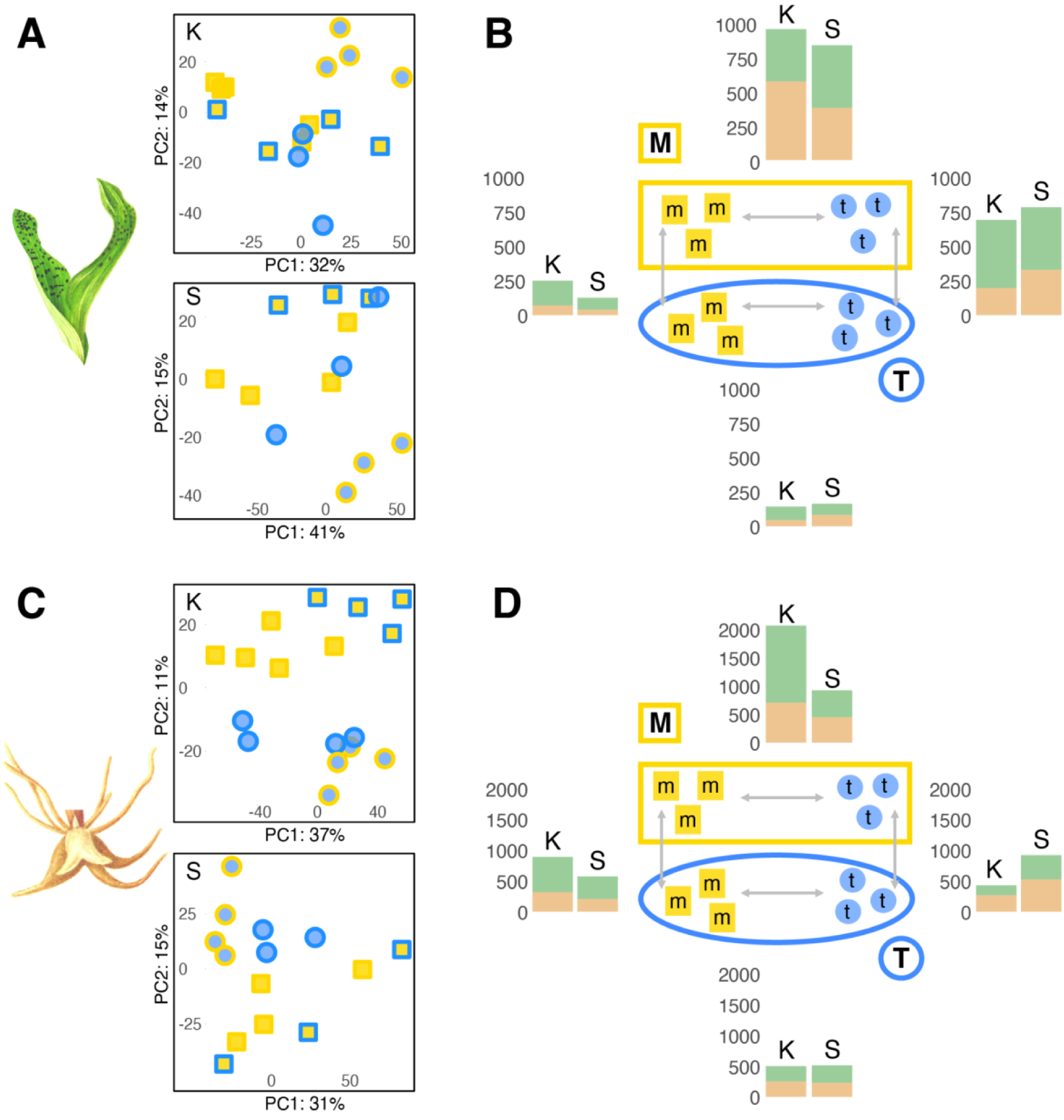
Variability of expression profiles of *Dactylorhiza majalis* and *D. traunsteineri* across locality, tissue and environment. **A, C** PCA plots for leaf and root data, respectively, separately per locality (K, Kitzbühel; S, St. Ulrich). Points represent biological replicates of *D. majalis* and *D. traunsteineri* (rectangles and circles respectively) grown in *D. majalis* or *D. traunsteineri* environments (symbols with blue or yellow lines, respectively). **B, D** Numbers of genes up- (green bars) or down-regulated (orange bars) between *D. majalis* (yellow symbols) and *D. traunsteineri* (blue symbols) and environments (yellow line, *D. majalis* environment; blue line, *D. traunsteineri* environment. B shows results from leaves, and D from roots. Left side bars show the number of DE genes at Kitzbühel (K), and the right side bars at St. Ulrich am Pillersee (S). The number of symbols in the experimental set up in the centre does not necessarily correspond to the actual analysed accessions per group.

Differential expression tests with DESeq2 v1.36.0 (Love et al., 2014) indicated that the two allopolyploids have more distinct profiles in the *D. majalis* environment (hereafter M environment), compared with the nutrient-poor habitat of *D. traunsteineri* (hereafter T environment), and this pattern was highly consistent between the localities and tissue types (Fig. 2B and D). In leaves and roots, respectively, 978 and 2,066 genes were differentially regulated between the allopolyploids in Kitzbuhel in the M environment, and 857 and 923 in St. Ulrich (Supplementary Table 2). However, only 143 and 501, and, respectively 163 and 512 genes were differentially expressed for the same comparisons in the T environment.

On the other hand, *D. traunsteineri* exhibited a greater number of DEGs in leaf tissue between native and non-native environments at both localities compared with *D. majalis* under the same conditions (Fig. 2B), however this trend was visible in root tissue only at St. Ulrich (Fig. 2D).

### Constitutive and plastic expression

Root tissue had more genes expressed (i.e., with at least one count per million in at least three individuals) overall compared with leaf tissue: 35,251 and 34,844 genes were expressed in root tissue in Kitzbuhel and St. Ulrich respectively, compared with 30,808 and 28,872 genes in leaf tissue for the same localities. Of these, only 30 and 29 genes in leaf tissue were constitutively differentially expressed between the two allopolyploids in Kitzbuhel and St. Ulrich respectively, and 51 and 63 genes in roots for the same localities (Fig. 3). All four comparisons showed a significantly greater number of constitutively expressed genes than expected by chance (p < 0.05, super exact test, Supplementary Tables 2-4). Electron transport during photosynthesis, pH regulation and transmembrane transport appear to be enriched among constitutively differentially expressed genes between the allopolyploids in leaves, whereas for the same comparison in roots, detoxification activity, carbohydrate transmembrane transport, and nucleotide and nucleoside metabolism is enriched (Fig. 4; Supplementary Table 5). Functional terms differentially regulated between the allopolyploids in the M environment only included cell wall biogenesis and defense response genes that are downregulated in the leaves of *D. traunsteineri*, and photosynthesis and signalling pathway that are upregulated in the same conditions. In turn, in the T environment, the leaves of *D. majalis* upregulate some photosynthesis genes. In roots, in the M environment several components of transmembrane transport differ between the two allopolyploids, whereas in the T environment apart from differences in transmembrane and water transport between the allopolyploids, some genes involved in response to heat and response to nitrogen compound are downregulated, and growth is upregulated in *D. majalis* (Fig. 4; Supplementary Table 5).

**Figure 3.**
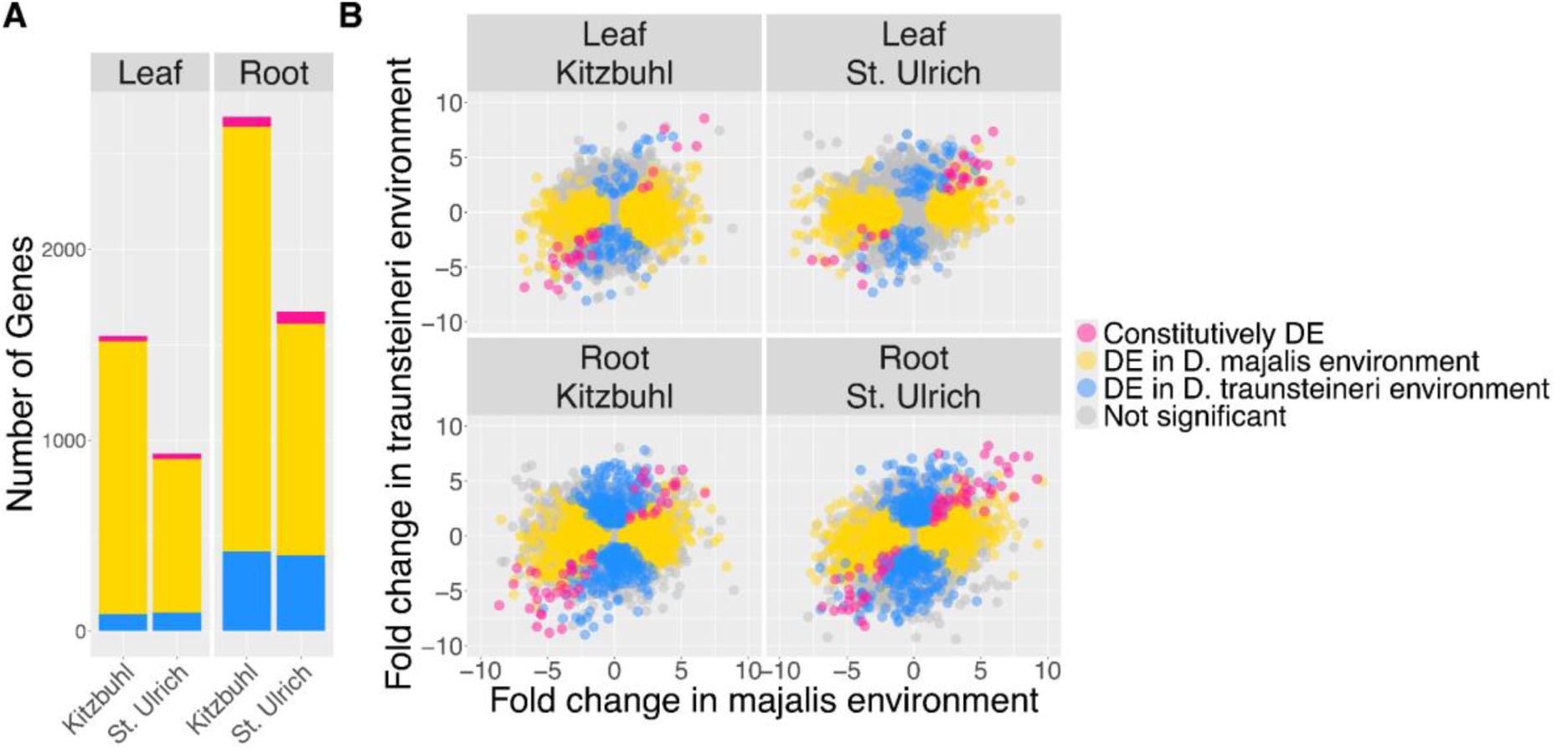
**A** Number of genes differentially expressed between *Dactylorhiza majalis* and *D. traunsteineri* in both environments (pink), in the *D. majalis* environment only (yellow), and in the *D. traunsteineri* environment only (blue). **B** Log fold change in gene expression between *Dactylorhiza majalis* and *D. traunsteineri* in the *D. traunsteineri* environment (Y-axis) and between *D. majalis* and *D. traunsteineri* in the *D. majalis* environment (X-axis). *D. majalis* was arbitrarily set as the base condition, therefore positive values indicate upregulation in *D. traunsteineri* relative to *D. majalis* and negative values indicate upregulation of *D. majalis* relative to *D. traunsteineri*. Genes shown in colour are designated differentially expressed (FDR < 0.05).

**Figure 4.**
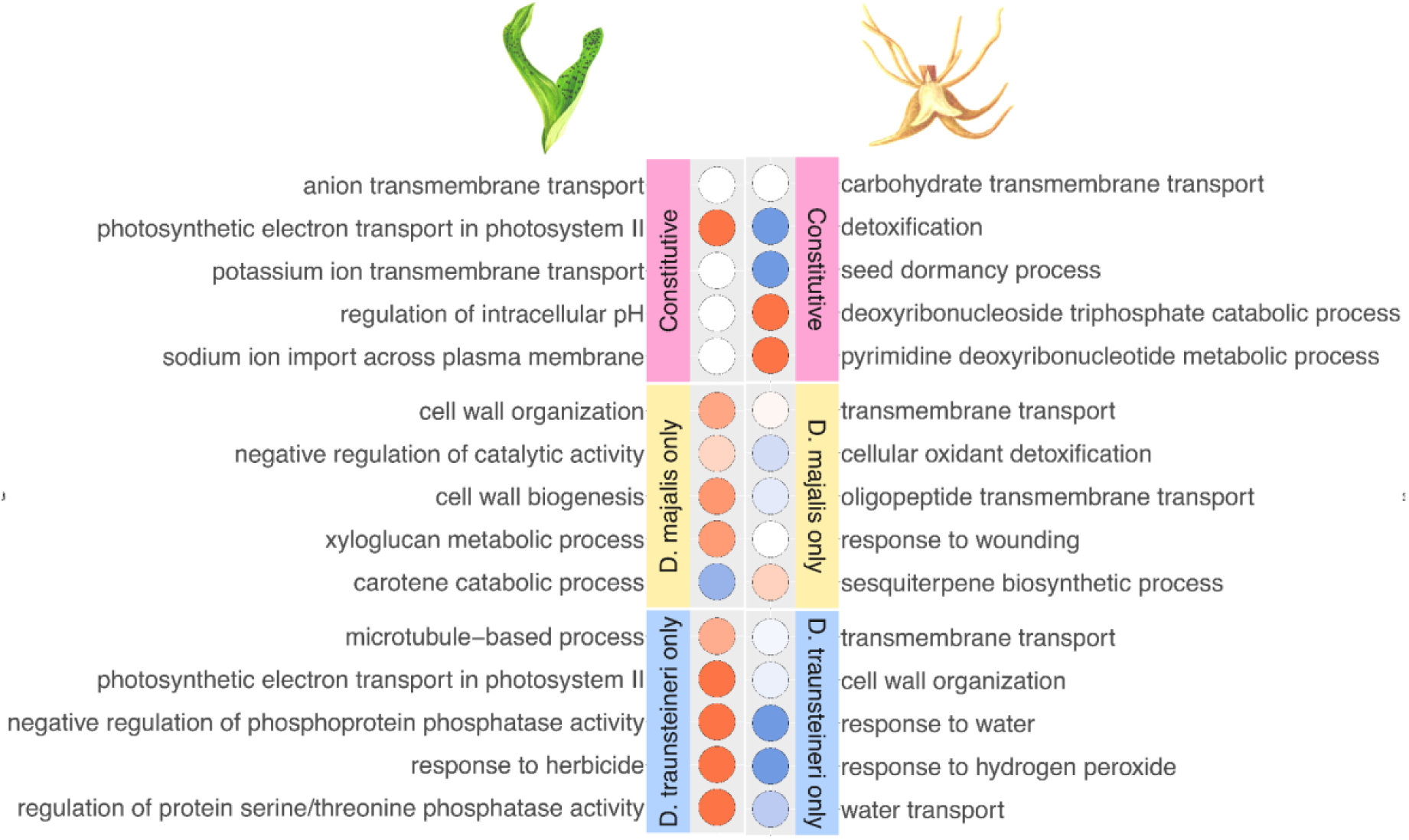
Top five GO terms significantly enriched (Fisher’s exact test p < 0.05) in genes that are differentially expressed between *Dactylorhiza majalis* and *D. traunsteineri* in both species’ environments (constitutive DEGs, pink panel), genes which are differentially expressed only in only the *D. majalis* environment (yellow panel) and genes which are differentially expressed only in the *D. traunsteineri* environment (blue panel). GO term categories are coloured by Z-score, indicating the directionality of gene expression underlying the enrichments, GO terms coloured in red contain more genes upregulated in *D. majalis*, GO terms coloured in blue contain more genes upregulated in *D. traunsteineri*. Results for leaf tissue samples are shown in the left panel, results for root tissue samples in the right panel. Differentially expressed genes for comparisons in Kitzbuhel and St. Ulrich are combined prior to functional enrichment analysis. Functional enrichment was performed on differentially expressed genes in each category (FDR < 0.05) using TopGO using the weight01 algorithm.

The environmental differences between the native habitats of *D. majalis* and *D. traunsteineri* elicit species specific plastic responses in either species when transplanted to the alternative environment (see *D. traunsteineri* plastic and *D. majalis* plastic, respectively, in Figure 1). In general, root tissues showed more transcriptional plasticity than leaves after transplantation of each species to the alternative environment (except *D. traunsteineri* in Kitzbühel), and both tissues tended to upregulate more genes than downregulate (except the roots of *D. traunsteineri*; Fig. 2B and D; Supplementary Table 2). The great majority of genes differentially regulated between environments were unique to a species and locality (Fig. 5) likely reflecting the divergent population histories across localities in particular for *D. traunsteineri* (see Supplementary Fig. 1, and Balao et al. 2016, Brandrud et al., 2020). *Dactylorhiza traunsteineri* leaf tissue showed the most overlap in plastic genes between Kitzbuhel and St. Ulrich of 17.6%, however most other comparisons showed fewer than 10% of plastic genes were replicated across both localities.

**Figure 5.**
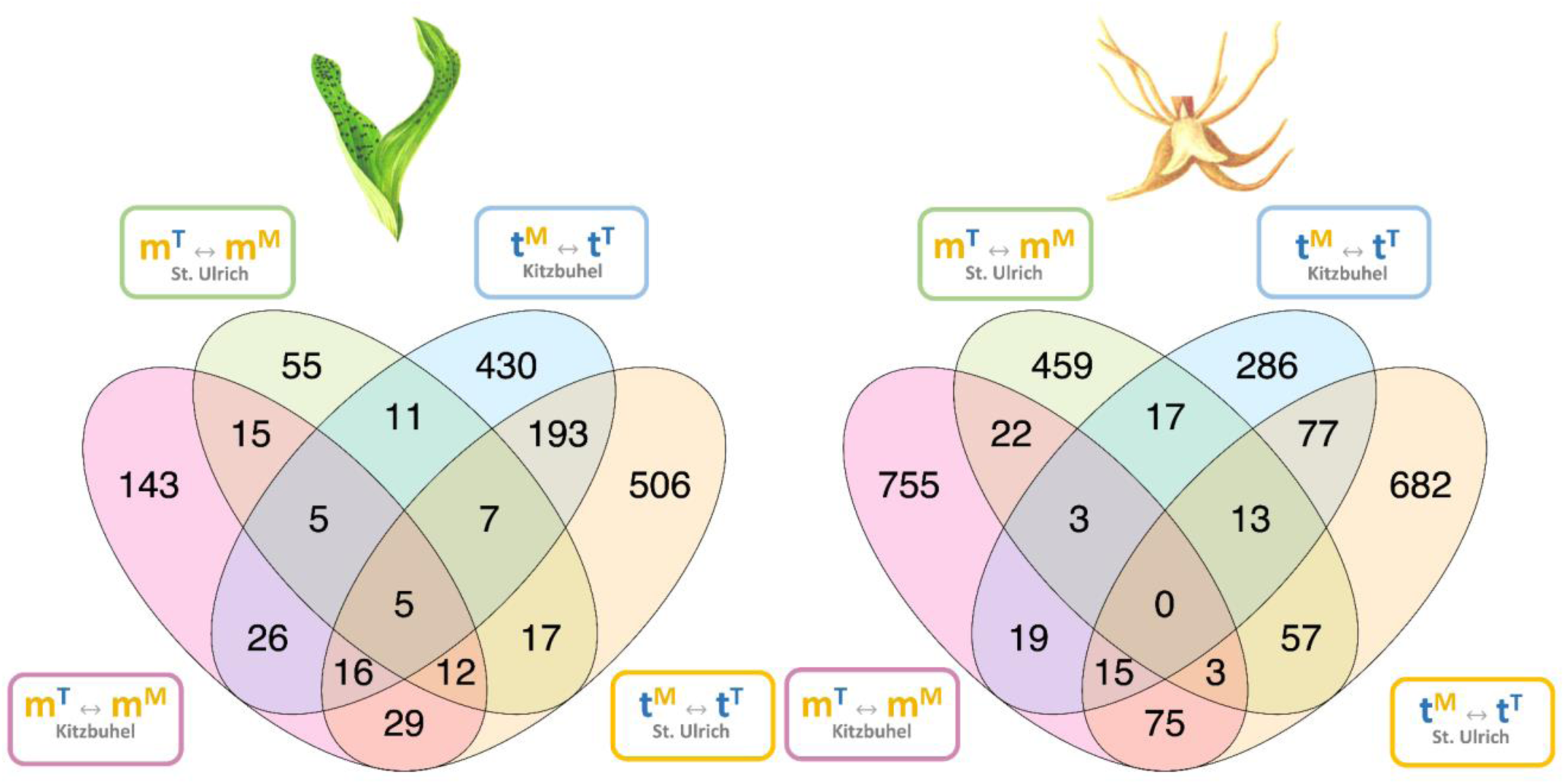
Venn diagram of plastic DE genes in each species (m, *D. majalis*; t, *D. traunsteineri*) when transplanted to the alternative environment (M, *D. majalis* environment; T, *D. traunsteineri* environment) (FDR < 0.05, log fold change > 1.5) within the same locality (K, Kitzbuhl; S, St. Ulrich). Labels follow Fig. 1b and 1d.

To better understand the functional context of plasticity in *D. majalis* and *D. traunsteineri*, we performed GO term enrichment analysis to identify significantly overrepresented functional categories (Fisher’s exact test p < 0.05) in genes exhibiting plasticity in their expression pattern (log fold change > 1.5, FDR < 0.05 in a single species grown in non-native versus native environment). To emphasise species-specific plastic responses, we focussed on functional categories which tended to be found disproportionately in either *D. majalis* or *D. traunsteineri,* calculating Z-score on proportion of GO terms from each species mapping to GOSlim terms and excluding functional categories with a Z-score of less than 0.4, and thus deemed to be found in roughly equal proportions in both species (see methods, for a full list of enriched functional terms see Supplementary Table 6). GOSlim terms disproportionately found to be enriched in *D. majalis* plastic gene expression were signal transduction, anatomical structure morphogenesis, cellular homeostasis and growth, and those in *D. traunsteineri* were DNA metabolic process, the response to abiotic stimulus (including light, ionizing radiation, heat and water deprivation), and photosynthesis. GO terms classified under these GOSlim terms are shown in Fig. 6.

**Figure 6.**
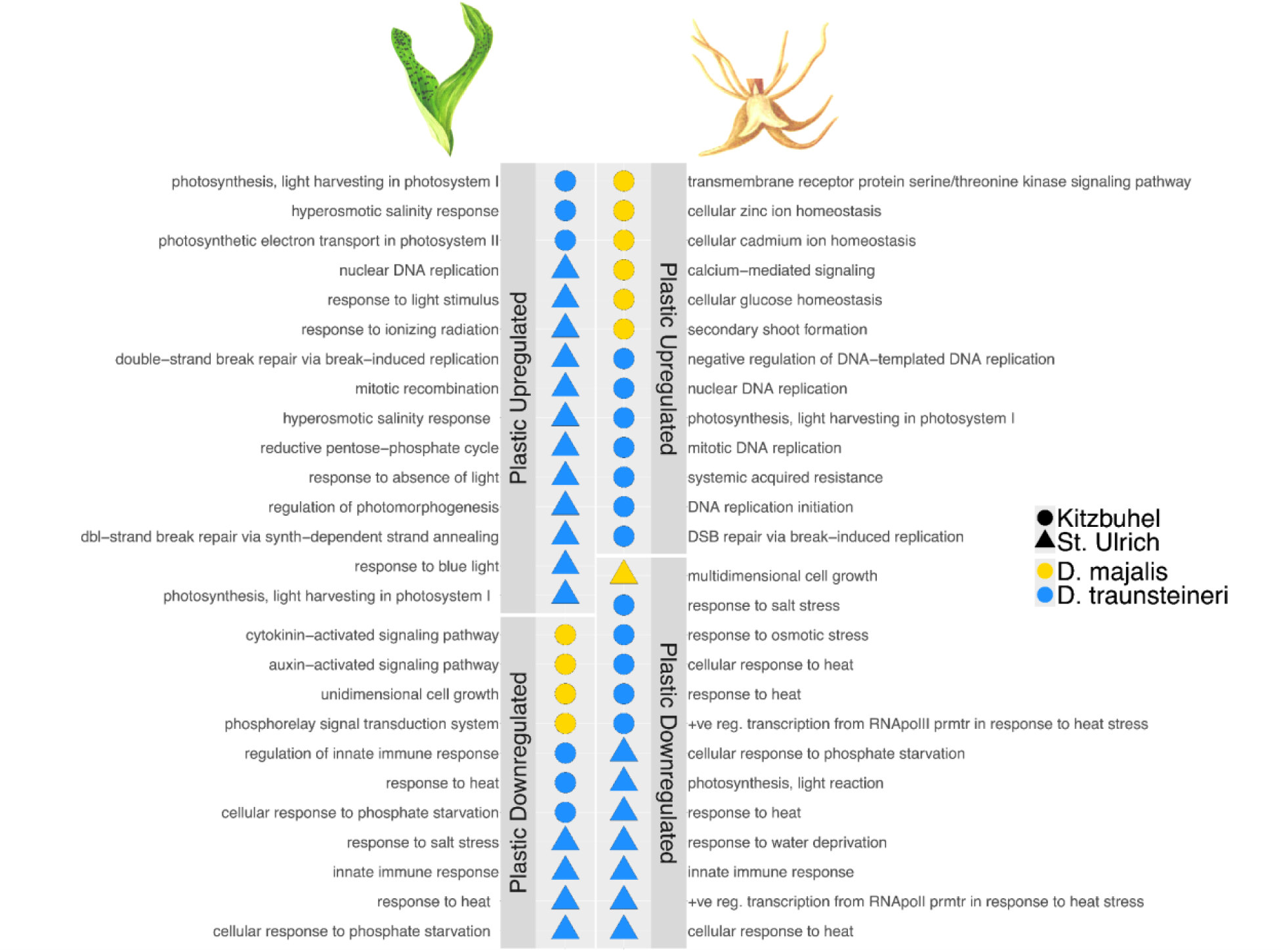
GO terms significantly enriched (Fisher’s exact test p < 0.05) in genes differentially expressed in each species upon transplantation to the environment of the opposite species, by tissue and locality. GO terms enriched in upregulated genes are shown in upper panels, and downregulated in lower panels. Only GO terms mapped in different proportions to respective GOSlim terms between *D. majalis* and *D. traunsteineri* are visualised, showing GO terms with a Z-score of > 0.4 (see Methods, for a full list of enriched GO terms see Supplementary Table 6).

Photosynthetic genes, including light response, and genes related to sensing are upregulated in *D. traunsteineri* leaves grown in the M environment, as were genes responding to hyperosmotic salinity stress and double strand break repair (Fig. 6). In turn, at both localities it consistently downregulated in the same conditions genes involved in response to heat, response to phosphate starvation and innate immune response. The leaves in *D. majalis* showed no enrichments, except for downregulated genes when grown in the T environment at Kitzbuhel, for which growth and regulation of growth through auxin and cytokinin activated pathways were overrepresented.

In root tissues, enrichments for *D. traunsteineri* upregulated genes were obtained only at Kitzbuhel, and they were related to mitosis and DNA replication, and systemic acquired resistance genes when grown in the M environment. *Dactylorhiza traunsteineri* roots further downregulated genes related to phosphate starvation, water deprivation, and salt and heat stress, as well as genes related to innate immune response. When grown in the T environment at Kitzbühel, *D. majalis* root tissue upregulated homeostasis of glucose and metal ions cadmium and zinc, the formation of secondary shoots, and genes related to the serine/threonine kinase transmembrane receptor pathway. *D. majalis* downregulated genes related to multidimensional cell growth in root tissue when grown in the T environment at St. Ulrich.

### Fungal taxonomic profiles of native and transplanted root samples

We sought to characterise the fungal species associated with the roots of *Dactylorhiza majalis* and *D. traunsteineri* because mycorrhizal and other fungi play an important role as determinants of fitness through orchid life cycles (Perotto & Balestrini, 2024; H. N. Rasmussen, 1990; H. N. Rasmussen & Rasmussen, 2009). The RNA-seq data used for this analysis was not collected for the sole purpose of fungal taxonomic profiling, and technical steps involved in data collection such as ribosomal sequence depletion may reduce the power of the analysis, but we argue that the standardisation of library preparation across samples will preserve inter-sample variation despite reduction of overall signal. Mapping RNA-seq reads to the Fungal RefSeq database with MiCoP (LaPierre et al., 2019) identified fungal community profiles which differed the most between the environments of *D. majalis* or *D. traunsteineri*, rather than by orchid species (Fig. 7).

**Figure 7.**
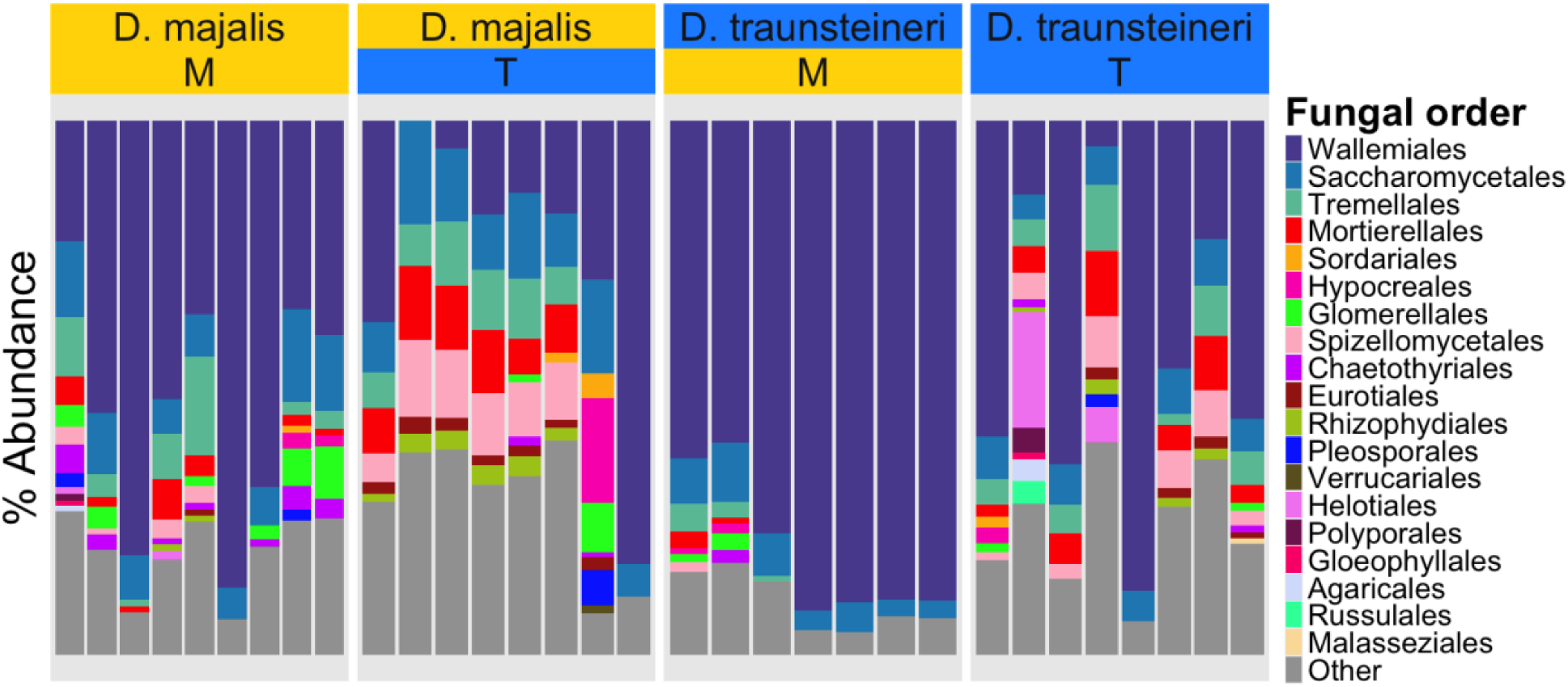
Percent abundance of fungal orders identified in root RNA-seq per sample. Fungal orders which were unclassified or present at a proportion of less than 1% are labelled “Other”. Transplantation of *D. majalis* and *D. traunsteineri* to the environment of the opposite species had the effect of shifting the fungal community profile of root tissue to that of the surrounding environment, rather than root tissue reflecting native fungal community profiles.

The M environment was characterised by the dominance of the order Wallemiales, a small group of xerophilic fungi in Basidiomycota (Zalar et al., 2005). Although this taxon was also often well represented in root tissues from the T environment, its lower incidence compared with the alternative environment was accompanied by a higher mean relative abundance of non-Wallemiales fungal taxa (46.8% and 19.7% mean relative abundance of non-Wallemiales fungal taxa in the M environment in *D. majalis* and *D. traunsteineri* respectively, compared with 74.4% and 55.7% in the T environment for the same species). The remaining taxonomic composition of the fungal community profiles included taxa which are known to be mycorrhizal partners to some orchids, such as Russulales (Looney et al., 2022), Mortierellales (Herrera-Rus et al., 2020) and Polyporales (Downing et al., 2020), as well as orders containing potentially pathogenic fungal species such as Glomerellales (Baroncelli et al., 2018).

Alpha diversity as measured by Shannon index (Fig. 8) showed a clear distinction in the diversity and evenness of fungal taxa in the M environment compared with the T environment, the latter showing near-consistently higher diversity for both species at both localities. Higher fungal diversity in the T environment compared with M was replicated across both Kitzbühel (M mean = 1.02, T mean=1.55) and St. Ulrich (M mean = 0.55, T mean=1.28). However, a statistically significant difference in Shannon Index was found only between individuals at M and T environments in St. Ulrich (Welch t-test, p=0.049), with no significant difference identified in Kitzbühel (Welch t-test, p=0.075).

**Figure 8.**
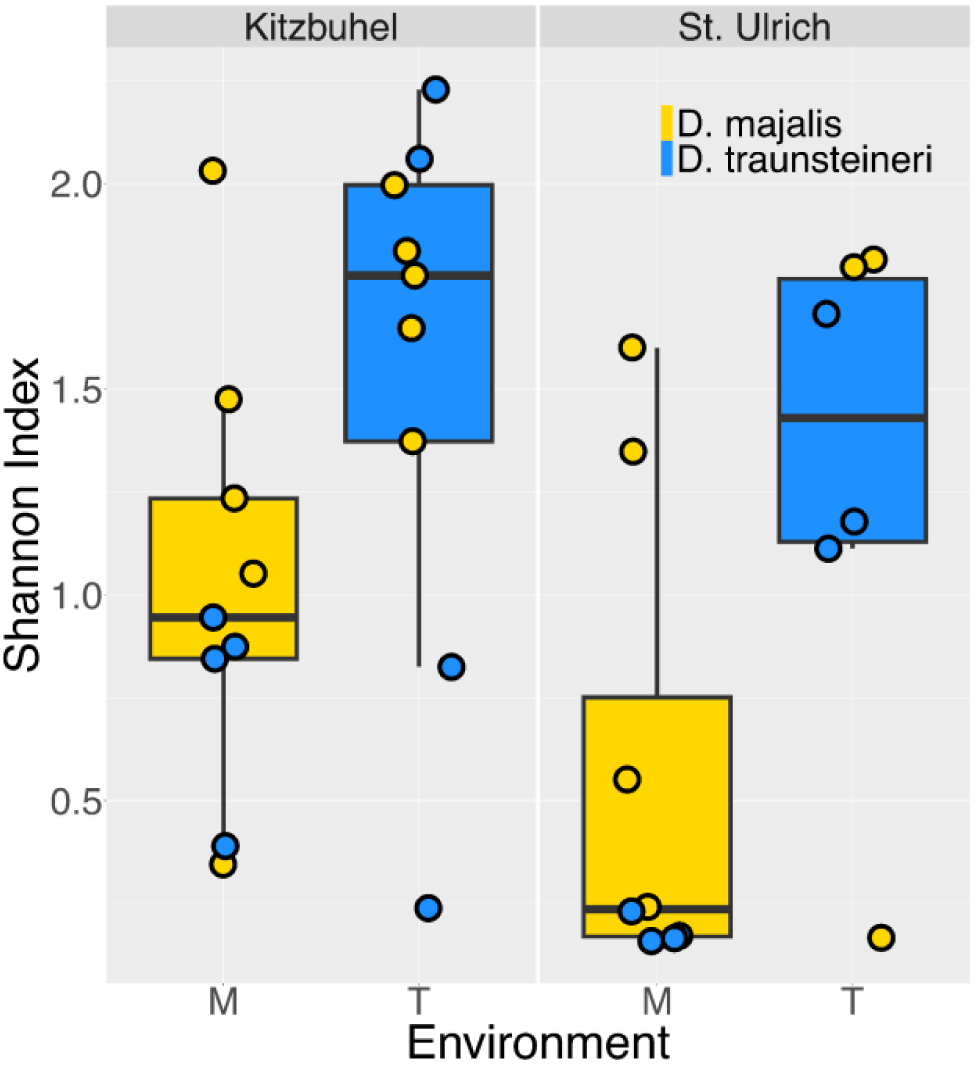
Alpha diversity of each environment (that of *Dactylorhiza majalis* or *D. traunsteineri*, denoted M and T respectively) and each locality (Kitzbuhel or St. Ulrich, left and right panels respectively) measured using Shannon Index. Shannon index was calculated using relative abundance of fungal order, and is shown across samples from both species per locality and environment as circled points (*D. majalis* in yellow and *D. traunsteineri* in blue).

LEfSe analysis (Supplementary Fig. 2) confirmed that Wallemiales was the only order significantly discriminative of the M environment (LEfSe LDA=3.14, p=0.0092). Wallemiales showed a higher mean relative abundance in the M environment, with between 49.4% and 90.3% mean relative abundance of Wallemiales regardless of species, compared to between 18.4% and 43.4% in the T environment. In contrast, three orders, Mortierellales (LEfSe LDA=2.44, p=0.0011), Spizellomycetales (LEfSe LDA=2.48, p=0.0010) and Tremellales (LEfSe LDA=2.21, p=0.022), were significantly discriminative of the T environment. The two most significantly discriminative orders of the T environment, Mortierellales and Spizellomycetales, had a mean relative abundance of between ca. 5 and 10% in *D. majalis* transplants and *D. traunsteineri* native individuals in both St. Ulrich and Kitzbühel. In contrast, these orders were found only in Kitzbühel in the M environment at a mean relative abundance of over 2%, with a maximum mean relative abundance of ca. 4%. Only one order (Tremellales, LEfSe LDA=2.32, p=0.0022) was found to be discriminative of the St. Ulrich and Kitzbuhel localities when testing for differential abundance using locality as the main class and environment as a subclass, suggesting that the M and T environments were the predominant drivers of fungal community differences and that this effect is relatively stable across the two localities.

Due to absence of usual orchid mycorrhizal taxa from the MiCoP database used for the above analyses, a closer inspection of previously reported key orchid fungal partners (Bateman et al., 2014; Jacquemyn et al., 2016; Read et al., 2024) was performed. Mapping of RNA-seq reads (see Methods) to *Ceratobasidium*, *Tulasnella*, *Serendipita*, *Sebacina* and *Russula* representative ITS sequences supported the presence of all tested taxa in both species and environments, *Ceratobasidium* being the most abundant taxon and *Russula* the least abundant. Variation in abundance of each fungal taxon was low (Supplementary figure 3, Supplementary figure 4) between species per treatment (e.g. *Dactylorhiza majalis* native vs. *D. traunsteineri* native) and within species between treatments (e.g. *D. majalis* native vs. *D. majalis* transplant), inter- and intra-specific comparisons tending to fall within the same range per fungal taxon.

## Discussion

Ecological plasticity is an important phenomenon that allows species to extend their genetically encoded trait optima, enabling them to better withstand environmental fluctuations (Fox et al., 2019) or extend their ranges (Richards et al., 2006). Polyploid plasticity is expected to be extensive, but it is often examined in the context of neopolyploid establishment, as it plays a critical role in overcoming obstacles such as reduced initial genetic diversity, minority cytotype exclusion and competition with diploids (Anneberg et al., 2023; Husband, 2000). Polyploids form recurrently, which enriches their genetic pool, but it can also lead to the development of distinct lineages, starting from different diploid populations, and each following independent evolutionary trajectories. The formation of stable polyploid species relies, among others, on the reinforcement of reproductive barriers to prevent genetic assimilation by more established cytotypes. However, adaptive introgression of genes can also increase the fitness of establishing polyploid taxa (Chapman & Abbott, 2010; Han et al., 2015). In this context, ecological plasticity can allow polyploids to colonise and establish themselves in novel environments, but also tolerate environments inhabited by related taxa where genetic exchange can occur, thus potentially exposing establishing polypoids to opposing evolutionary forces.

### Dominant plastic responses are punctuated by fixed transcriptional differences between species

To explore the further implications of ecological plasticity for repeated polyploid formation, we conducted reciprocal transplantations with *D. majalis* and *D. traunsteineri.* These sibling allopolyploids occupy distinct habitats (meadows with mesic soils versus low nutrient fens), an ecological divergence that has been previously shown to be associated with distinct features of leaf chemistry, light harvesting, photoprotection, nutrient transport and stomatal activity between the two allopolyploids (Wolfe et al., 2023). Our goal here was to quantify the plastic expression range of each species relative to the amount of fixed expression differences accumulated between them. Due to the lack of a reference genome for the maternal diploid *D. fuchsii*, and the relatively low divergence in coding regions between the two parents (Balao et al., 2017), our analyses focused on gene level expression and did not attempt to quantify plasticity at the homoeolog level. We found very few constitutive DEGs, with more such genes identified in roots compared with leaves, and this consistently among the two localities investigated. The higher number of constitutive DEGs in roots aligns with the widespread differences in soil nutrient profiles between the specific environments of the two allopolyploids (Fig. 1; see also Paun et al., 2011; Wolfe et al., 2023), suggesting that the below-ground environment exerts greater divergent selective pressure on *D. majalis* and *D. traunsteineri*.

Rather than finding many constitutive DEGs, most DEGs between *D. majalis* and *D. traunsteineri* were environment-specific (Fig. 3). Notably, many more DEGs between the allotetraploids were found in the M environment than in the T. Our results further show that *D. traunsteineri* undergoes in general greater transcriptional remodelling in the alternative environment, than the vice versa situation. This explains at least in part the generally observed asymmetric gene flow that is more intense from *D. majalis* to *D. traunsteineri* (Balao et al., 2016; Brandrud et al., 2020). More widespread and competitive species like *D. majalis* (Paun et al., 2011) are thought to have higher levels of plasticity (Hiatt & Flory, 2020) than those with narrow ecological preferences such as *D. traunsteineri*. The transplantation of *D. majalis* to the T environment may fall more within its tolerated range, and hence require fewer changes. Alternatively or in addition, as the oldest of the two allopolyploids, *D. majalis* may be further along the transcriptional diploidization process.

Adaptation to the nutritionally depauperate environment of *D. traunsteineri* requires significant metabolic modifications (Wolfe et al., 2023). If the basis for these modifications is rather plastic, the transplantation of *D. traunsteineri* to the *D. majalis* environment may allow it to return to a transcriptional baseline to make use of the newly available resources, a possibility that is reflected in the functions of plastic genes (Fig. 6 and discussed later). Plasticity can mitigate unfavourable interspecific interactions (Otte et al., 2016), providing a buffer that prevents competitive exclusion or genetic swamping, thereby facilitating species coexistence (Hess et al., 2022). Polyploid plasticity, especially along environmental clines, is important for achieving the ecological speciation required for successful polyploid establishment (Shimizu-Inatsugi et al., 2017). The heterogeneous soil types occupied by *D. majalis* and *D. traunsteineri* likely served as the foundation for the ecological divergence between them.

Transcriptional plasticity allowing *D. traunsteineri* to survive in nutritionally poor environments, characterised by low competition, may be rewarded by allowing it to successfully establish as a young allopolyploid, developing genetic isolation away from the swamping effects of hybridisation when grown in sympatry with its siblings and diploid parents. Whereas global transcriptional profiles do not always clearly discriminate *D. majalis* and *D. traunsteineri* (Fig. 2A and 2C), the small number of constitutive DEGs may represent evidence of nascent differentiation in response to habitat-specific conditions.

### Sibling allopolyploids plastically alter nutrient acquisition, photosynthetic capacity and osmotic regulation in the alternative habitat

Growing each sibling allopolyploid in the habitat of the other stimulated remodelling of functions reflecting their previously described habitat differences such as soil chemistry and nutrient availability (Wolfe et al., 2023); *D. traunsteineri* downregulated genes concerned with cellular response to phosphate starvation in leaves and roots, suggesting an alleviation of pressure in nutrient availability when grown in the *D. majalis* environment.

While upregulation of nutrient starvation functional terms is not seen in *D. majalis*, terms involving root system architecture remodelling and zinc homeostasis are upregulated in *D. majalis* root tissue (Supplementary Table 6, Fig. 6), reflecting a suite of processes responsive to nutrient starvation and abiotic stress (Bechtaoui et al., 2021; Shao et al., 2020). Root architecture remodelling is used by many angiosperms to modulate rate of water and nutrient absorption by increasing root surface area, often through increasing the density of root hairs (Kiba & Krapp, 2016; Sánchez-Calderón et al., 2005). Orchids have little to no root hairs present in non-tuberous roots (Zhang et al., 2018), however orchid roots have been shown to be highly plastic to different growing conditions (de Lima & Moreira, 2022) and structural changes are likely induced in response to a change in nutrient availability. Phosphate limitation has been shown to be a key modulator of zinc bioavailability, forming a complex homeostatic interaction in plant responses to nutrient deficiency (Bouain et al., 2014; Lay-Pruitt et al., 2022).

Consistent with differences in water availability between the environments of *D. majalis* and *D. traunsteineri,* the latter shows a broad response to osmotic pressure, downregulating genes involved with water deprivation and salt stress, and upregulating genes annotated to hyperosmotic salinity response. Osmotic regulation is required to maintain efficient stomatal conductance (Bandurska, 2022) and is an important modulator of root system depth and breadth (Deak & Malamy, 2005), and the preponderance of *D. traunsteineri* plastic genes annotated to such terms may reflect the extent of the osmotic remodelling usually required in its native environment. To a lesser extent, *D. majalis* root tissue showed signs of plastically adjusting to an osmotically altered environment, upregulating important abiotic stress sensitive functions such as calcium-mediated signalling (Wilkins et al., 2016), as well as functions modulating glucose homeostasis, a key osmoprotectant (Ahmad et al., 2020). Wolfe et al. (2023) showed a lower energetic efficiency of photosynthesis in *D. traunsteineri* compared with *D. majalis*, and a higher rate of photooxidative regulation through quenching. Leaf tissue of *D. traunsteineri* grown in the *D. majalis* environment shows plastically reduced levels of gene expression in pathways known to alleviate oxidative stress and maintain efficient photosynthesis such as glutathione (Müller-Schüssele et al., 2020) and L-ascorbate (Chen et al., 2021) (Supplementary Table 6). Moreover, plastically upregulated genes are enriched for photsynthetic and photomorphogenic functions, indicating a plastic increase in *D. traunsteineri* photosynthetic efficiency upon release from pressures in its native environment.

Interestingly, up- and downregulated genes in *D. traunsteineri* root tissues were enriched in some photosynthesis related terms. Photosynthesis tends to be inhibited under low nutrient availability (Farhat et al., 2016; Mu & Chen, 2021), however due to roots being non-photosynthetic tissue, study of photosynthesis regulation in root tissue is sparse. Kang et al. (2014) identified photosynthetic genes downregulated in the root tissue of *Arabidopsis thaliana* during phosphorous starvation, speculating that phosphorous depletion may stimulate aberrant photosynthesis to produce elevated levels of reactive oxygen species (ROS). Both nutrient deficiency (Ahanger et al., 2017; Kim et al., 2010) and photosystem I of the photosynthetic apparatus (Furutani et al., 2020) have been linked to elevated levels of ROS. The response to ROS appears to be diminished in *D. traunsteineri* when grown in the non-native *D. majalis* environment, as indicated by the downregulation of genes enriched in L-ascorbic acid and glutathione pathways, known for their role in ROS scavenging (Akram et al., 2017).

### Interspecific fungal community differences

Whereas fungal reads were extracted from the *Dactylorhiza* root RNA-seq data, it is not clear what proportion of the sequence reads originate from genuine orchid mycorrhizal partners rather than bystander fungi present in the surrounding rhizosphere. Nevertheless, we find a consistent difference in the mycobiome associated with the sibling allopolyploids; diversity measures show higher alpha diversity in the *D. traunsteineri* environment (Fig. 8) regardless of species or locality compared with the *D. majalis* environment, which was dominated by Wallemiales taxa and exhibited smaller proportions of taxa well represented in the *D. traunsteineri* environment such as Spizellomycetales and Mortierellales. Three orders were discriminant of the *D. traunsteineri* environment (Mortierellales, Spizellomycetales and Tremellales), whereas Wallemiales was the only fungal order which was significantly discriminant of the *D. majalis* environment. This result is interesting given the limitation of resources such as nitrogen and phosphorus in the *D. traunsteineri* environment, which may prompt adapted plants to cultivate relationships with a more diverse set of specialist mycorrhizal partners to alleviate such nutrient deficiencies (Kohler et al., 2015; Wyatt et al., 2014).

Absence of key mycorrhizal species in the RefSeq database used by MiCoP resulted in these important taxa being absent from our characterization of mycobiota. We were able map some reads from root RNA-seq datasets to representative ITS sequences of previously identified orchid fungal partners, confirming their presence in *Dactylorhiza majalis* and *D. traunsteineri* root samples. Supporting previous studies (Jacquemyn et al., 2016; Pecoraro et al., 2018), *Ceratobasidium*, *Tulasnella* and *Sebacina* were the most abundant taxa of those tested (Supplementary figure 3), as was *Serendipita*, another taxon within the Sebacinales which has been previously shown to form associations with orchid roots (Fritsche et al., 2021). Interestingly, very low abundances of *Russula* were found in all *Dactylorhiza* samples of this study, despite extensive evidence of mycorrhizal association of this taxon with the roots of orchids (Dearnaley, 2007; Okayama et al., 2012). Leveraging of plant RNA-seq datasets to test for associated micro- and mycobiota extends the utility of such experiments and expands the scope of ecological studies in a cost-effective manner. Using full genome fungal databases, approaches that use RNA-seq reads as input maximise read usage and circumvent problems introduced by standard ribosomal depletion steps during library preparation that preclude the reliable use of ITS sequences for quantitative studies. Despite the clear benefits, fungal database annotation and curation is comparatively neglected, resulting in many fungal sequences remaining unannotated past the kingdom level and therefore not contributing to meaningful taxonomic research (Abarenkov et al., 2022). Nevertheless, our ability to detect common orchid fungal symbionts by mapping RNA-seq reads to selected ITS sequences highlights the potential of such approaches for basic presence-absence analyses and merits further investigation.

The association of orchid species with mycorrhizal fungi represents a fascinating and well studied partnership, and the broader role of fungal associations in modulating plant health represents an important component in understanding plant-environment dynamics. Moreover, the correlation of fungal community profiles with ecologically divergent plant groups suggests that symbiont recruitment and selection by plants may be itself plastic (i.e., environment-dependent) and reflective of the environmental challenges plants face (Davison et al., 2020). The environments of *D. majalis* and *D. traunsteineri* differ markedly in their nutrient availability, and the differences in fungal community diversity could reflect previous studies showing resource limitation as a driver of locally adapted mycorrhizal associations (Johnson et al., 2010); however further studies are required to confirm whether this is the case in *Dactylorhiza*. Climatic niches for *D. traunsteineri* and *D. majalis* were also found to differ particularly according to temperature seasonality and mean temperature of the driest quarter (Wolfe et al., 2023), and previous studies of *D. majalis* have found symbiotic germination of seeds to be highly temperature dependent (H. Rasmussen et al., 1990), suggesting factors affecting germination success may also differ between *D. majalis* and *D. traunsteineri*. The study presented here transplanted adult plants to the environment of the other species, and consequently the germination phase of the life cycle was not captured. Further studies are needed to conclusively determine whether fungal partners play a role in the delimitation of *D. majalis* and *D. traunsteineri* native ranges, and more broadly whether such interactions help drive ecological divergence in nascent species.

### Conclusions

Hybridization and polyploidy are rich sources of biodiversity in the genus *Dactylorhiza*, but consequences of these processes can have contradictory effects. We show that divergent ecological preferences in nascent species *D. majalis* and *D. traunsteineri* are circumvented by transcriptomic plasticity, impeding the build-up of fixed transcriptomic differences. We further found that the nutritionally depauperate soil of *D. traunsteineri* harbours a more diverse community of fungal taxa than the nutritionally abundant soil of *D. majalis*. Taken with the broader role the rhizosphere plays in orchid germination and nutrient acquisition, we argue that divergent symbiotic interactions at key life stages may play an important role in the distribution of this pair of sibling *Dactylorhiza* species, and merits further investigation.

## Methods

### Study sites

The first locality selected for the transplantation experiments is near Lake Schwarzsee, north-west of Kitzbühel (Tyrol, Austria), which is the type locality of *D. traunsteineri* (Foley, 1990). In this area, the populations of *D. majalis* and *D. traunsteineri* are separated by a region lacking *Dactylorhiza* orchids, with the shortest distance between them being approximately 100m. Here, *D. majalis* grows in shady places under trees. *Dactylorhiza traunsteineri* plants are found in the open marshland closer to the lake, primarily on elevated and thus dryer locations (hummocks), nevertheless soil humidity is higher than in the *D. majalis* habitat. In general, the soil specific for *D. majalis* populations is significantly richer in most macro- and micronutrients as compared with *D. traunsteineri* (Wolfe et al., 2023). However, at the specific site in Kitzbuhel, the soil shows the opposite trend for phosphorus content (Fig. 1b).

The second locality is situated south of Lake Pillersee, northwest of St. Ulrich am Pillersee (Tyrol, Austria). Here the two species are also separated by an area (shortest distance ca 200m) that lacks either species, but it hosts rare individuals of the diploid parent *D. incarnata*. Within this area, the shortest distance between the two allopolyploids includes a ca 50m wide tree stand. Both species at St Ulrich am Pillersee occupy very humid soils, but the soil chemistry at this site agrees with the characteristic difference observed across their range (Fig. 1b). Previous genetic analyses have shown that gene flow between the two species is very prominent at Kitzbuhel, whereas admixture is also present to some extent at St. Ulrich am Pillersee, at least partly via backcrosses with diploid *D. incarnata* (Balao et al., 2016; Brandrud et al., 2020).

### Reciprocal transplantation and RNA sequencing

Ten wild individuals each of *D. majalis* and *D. traunsteineri* were randomly selected at each of the two localities. *Dactylorhiza* plants are perennials and grow a new tuber each growing season to support growth in the following season (Wolfe et al., 2023), therefore plants were sampled after two growing seasons to eliminate as much as possible the effects of the original environment. Tissue types used in this study were leaf and root; for leaf tissue the second mature leaf was sampled from each plant, and for root tissue horizontally radiating roots were sampled, distinguishing from root tubers which grow vertically downwards. Hereafter tissue types sampled will be referred to as ‘leaf’ and ‘root’ for brevity. The plants at Kitzbühel and St. Ulrich were sampled in 2018, on May 25 and 26, respectively. Leaf and root tissues were collected at each locality in the morning and stored in RNAlater (Sigma-Aldrich) at 4°C overnight, before transferring to −80°C. At collection, the transplanted plants were entirely removed from the sites.

Total RNA was extracted using the mirVana Isolation Kit (Thermo Fisher Scientific) according to manufacturer’s instructions, and quantitated using a NanoDrop ND-1000 Spectrophotometer (Thermo Scientific), using the wavelength ratio of A260/280 to estimate purity. A Plant RNA Pico kit was used on a 2100 Bioanalyzer (Agilent Technologies) to confirm the quality of the RNA isolates. DNA digestion was performed with DNase I (Promega), to remove any DNA contamination. Ribosomal RNA was depleted using the RiboZero Plant Kit for RNA-Seq (Illumina) according to manufacturer’s instructions. RNA was fragmented by hydrolysis for 2 minutes at 94° and cDNA synthesis was performed with random hexamers following the TruSeq Stranded RNA kit protocol (Illumina). To achieve strand specificity, the second strand was synthesised with dUTPs. The ribosomal depleted total RNA libraries were sequenced using directional pair-end sequencing on an Illumina NovaSeq instrument at the Next Generation Sequencing Facility of the Vienna BioCenter Core facilities (VBCF, www.viennabiocenter.org/vbcf). The RNA-seq data are available from GenBank BioProject PRJNA1151607.

Three accessions did not survive during the experiment, whereas two accessions were genetically identified as heavily introgressed between the two sibling allopolyploids and one as backcrossed towards the diploid *D. incarnata* (the latter at the locality of St. Ulrich). Finally, few accessions have been excluded either because of insufficient quality of the RNA (i.e., RNA integrity number was consistently lower than six across multiple RNA extractions) or of the sequence data obtained (i.e., showing prominently more genes with zero counts than the rest of the accessions). Across both allopolyploids, 16 leaf and 17 root samples from Kitzbuhel have been retained for the bioinformatic analyses, and 13 leaf and 14 root samples from St Ulrich (Supplementary Table 1).

### Genetic clustering

Quality filtered and adapter trimmed reads were mapped to the *D. incarnata* v.1.0 reference genome (Wolfe et al., 2023) using its .gff annotations and STAR v2.7.11a (Dobin et al., 2013) in two-pass mode, excluding any reads which mapped to more than ten positions. To investigate the genetic structure of accessions, we used ANGSD v0.941 (Korneliussen et al., 2014) for estimating genotype likelihoods, as this approach is suitable for RNA-seq data (Szukala et al., 2023), which is expected to include highly variable sequencing depth among genes and exons. For this purpose, the mapped bam files of the two tissues for each accession were merged with SAMtools v1.7 (Li et al., 2009). ANGSD was further run based on a SAMtools model (-GL 2), inferring the major/minor alleles (-doMajorMinor 1) and estimating allele frequencies (-doMaf 1). In the first round we retained SNPs that were identified outside annotated repeat regions, and which had a p-value lower than 2e-6 (-SNP_pval 2e-6), a minimum mapping quality of 20 (-minMapQ 20), a minimum base quality of 20 (-minQ 20), with data for at least 80% of individuals (-minInd 26) and a minor allele frequency corresponding to two accessions (-minMaf 0.06). In a second run we retained from the resulting variants only one per 2-kb window, aiming to reduce linkage disequilibrium between the markers. Covariance matrices computed from the obtained genotype likelihoods were used for principal components analyses (PCA) using pcANGSD v0.99 (Meisner & Albrechtsen 2018) and were plotted in R.

### Differential expression analysis

Following Wolfe et al. (2023) we quantified differential gene expression at the gene level, using reads mapped to the available *D. incarnata* genome. Uniquely mapped reads were summarised per tissue type across the coding sequence, excluding introns, for each sample using featureCounts from Rsubread v2.0.4 (Liao et al., 2019). Prior to differential expression analysis, the counts matrix was filtered to exclude low expression genes, retaining only those expressed at greater than one count per million in at least three samples, corresponding to the number of retained biological replicates in the smallest group in the analyses. DESeq2 v1.36.0 (Love et al., 2014) was used to identify differentially expressed genes (DEGs), normalising counts using median of ratios. To avoid incorporating signals of population structure and history, DEGs were detected for each locality (St Ulrich or Kitzbuhel) separately. Each test was performed to compare either the same species in different environments, or the two species in the same environment.

First, we identified constitutive gene expression differences, defined as DEGs between the sibling allopolyploids consistently identified in either environment, by finding genes which are significantly (p < 0.05, LogFC > 1.5) up- or downregulated between species in both environments; i.e. the intersection of DEGs found in comparisons m^M^ vs t^M^ and m^T^ vs t^T^ (where m stands for *D. majalis*, t for *D. traunsteineri*, superscript M for the *D. majalis* environment and superscript T for that of *D. traunsteineri*; see also “constitutive DE” comparison in Fig. 1c). To determine whether there is a significant overlap of DEGs between each comparison (i.e. whether we find more constitutive DEG than we expect by chance) we use the super exact test implemented in the SuperExactTest package (Wang et al., 2015) in R, using the number of genes included in the DESeq2 analyses as the background set from which constitutively DEGs are sampled.

Then we identified genes which showed plastic expression patterns (significantly differentially expressed, p < 0.05, LogFC > 1.5) in each species and locality when transplanted to the environment of the opposite species, by quantifying DEGs between t^M^ and t^T^ for *D. traunsteineri*, and between m^T^ and m^M^ for *D. majalis* (see “*D. traunsteineri* plastic” and “*D. majalis* plastic” comparisons respectively in Fig. 1c). Functional enrichment analyses were performed on the resultant groups of genes using the Bioconductor package topGO v2.48.0 (Alexa & Rahnenfuhrer, 2023) using the weight01 algorithm and all genes in the reference genome as background set.

To focus on species-specific patterns of constitutive and plastic gene expression, Z-score was calculated for significantly enriched functional terms to indicate the difference in number of genes mapped to GO terms or GO terms mapped to GOSlim terms between *D. majalis* and *D. traunsteineri*, a higher absolute number reflecting greater directionality towards one species or another. Z-score was calculated for number of genes mapping to GO terms as (#genes^majalis^ -#genes^traunsteineri^) / (genes^majalis^ + genes^traunsteineri^), and for proportion of GO terms mapping to GO slim terms as (%GOterms^majalis^ -%GOterms^traunsteineri^) / (%GOterms^majalis^ +%GOterms^traunsteineri^).

### Fungal taxonomic abundance analysis

We performed community profiling of fungal taxa using the RNA-seq root samples of *D. majalis* and *D. traunsteineri* plants grown in the two environments using MiCoP (LaPierre et al., 2019), a mapping based approach for classification and abundance of viral and fungal sequences. MiCoP uses a rapid read mapping approach to achieve more granular classification estimates which are more sensitive to low-abundance species than comparable methods such as kmers or megablast (Altschul et al., 1990). Paired end reads from each root sample were mapped with bwa mem v0.7.11 (Li et al., 2009) to the fungal RefSeq database accessed 7th November 2017 (Robbertse & Tatusova, 2011) included with the MiCoP release, followed by computation of fungal abundances from the alignment file. Quantification of diversity was performed with Shannon Index in the R package vegan v2.6.4 (Oksanen et al., 2009) calculating the Shannon index on relative abundance of fungal orders identified in each sample. Statistically significant differences in the Shannon diversity indices of the *D. majalis* and *D. traunsteineri* environments was tested per locality using a Welch Two Sample T-Test. Differential association of specific fungal taxa with the environment of either *D. majalis* or *D. traunsteineri* was analysed using LDA effect size (LEfSe, Segata et al., 2011). Features were scaled to sum to 10,000, using a p-value of 0.05 for both Kruskal-Wallis and Wilcoxon tests, and an effect size of discriminative features threshold of LDA > 3.

Finally, due to the absence of orchid mychorrizal taxa from the MiCoP database, the key fungal orchid partners derived from studies primarily located in Europe were identified and representative ITS sequences were downloaded from the UNITE database (Nilsson et al., 2019), selecting the following taxa and UNITE sequence ID for further inspection; *Ceratobasidium* AB520309, *Tulasnella* OP537806, *Serendipita* KF646113, *Sebacina* UDB000773 and *Russula* UDB05536919. RNA-seq reads from root samples were mapped against each representative sequence using bwa mem v0.7.11 (Li et al., 2009) and summarised using featureCounts from Rsubread v2.0.4 (Liao et al., 2019). Counts were FPKM normalised and visualised using the pheatmap package in R (Kolde, 2018).

## Acknowledgements

We thank Juliane Baar, Richard Bateman, Marie Brandrud, Mark Chase, Marcel Hirsch, Marie Huber and Daniela Paun for their contribution to the ideas and results presented here. Sequencing was performed at the Vienna BioCenter Core Facilities (VBCF; https://www.viennabiocenter.org/). Computational resources were provided by the Life Science Compute Cluster (LiSC) of the University of Vienna. Regional county administrations in Austria are acknowledged for issuing necessary permits.

## Data availability statement

The raw Illumina sequencing data will be deposited on NCBI SRA (GenBank BioProject PRJNA1151607) before the paper will be accepted for publication. The table of counts and the bioinformatics scripts are available from GitHub (https://github.com/katieemelianova/dactylorhiza/blob/main/dactylorhiza_DE.R).

## Supplementary figures

**Supplementary Figure 1.**
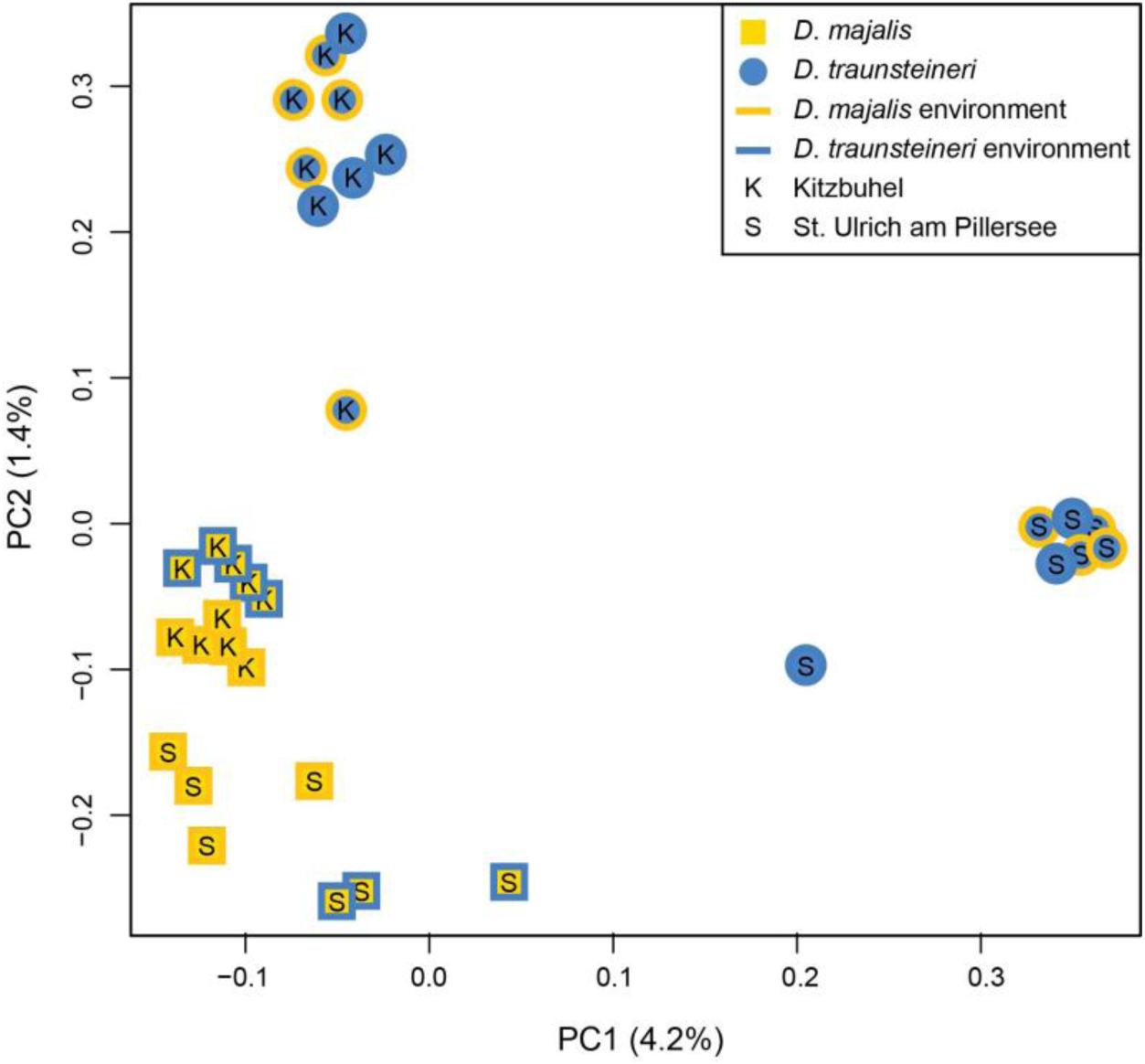
PCA plot based on genotypic variability expressed as genotype likelihoods in *Dactylorhiza majalis* and *D. traunsteineri* in the two localities tested (Kitzbühel and St. Ulrich am Pillersee). Symbols represent replicates of *D. majalis* (squares) and *D. traunsteineri* (circles) in native and transplanted habitats (according to the legend).

**Supplementary Figure 2.**
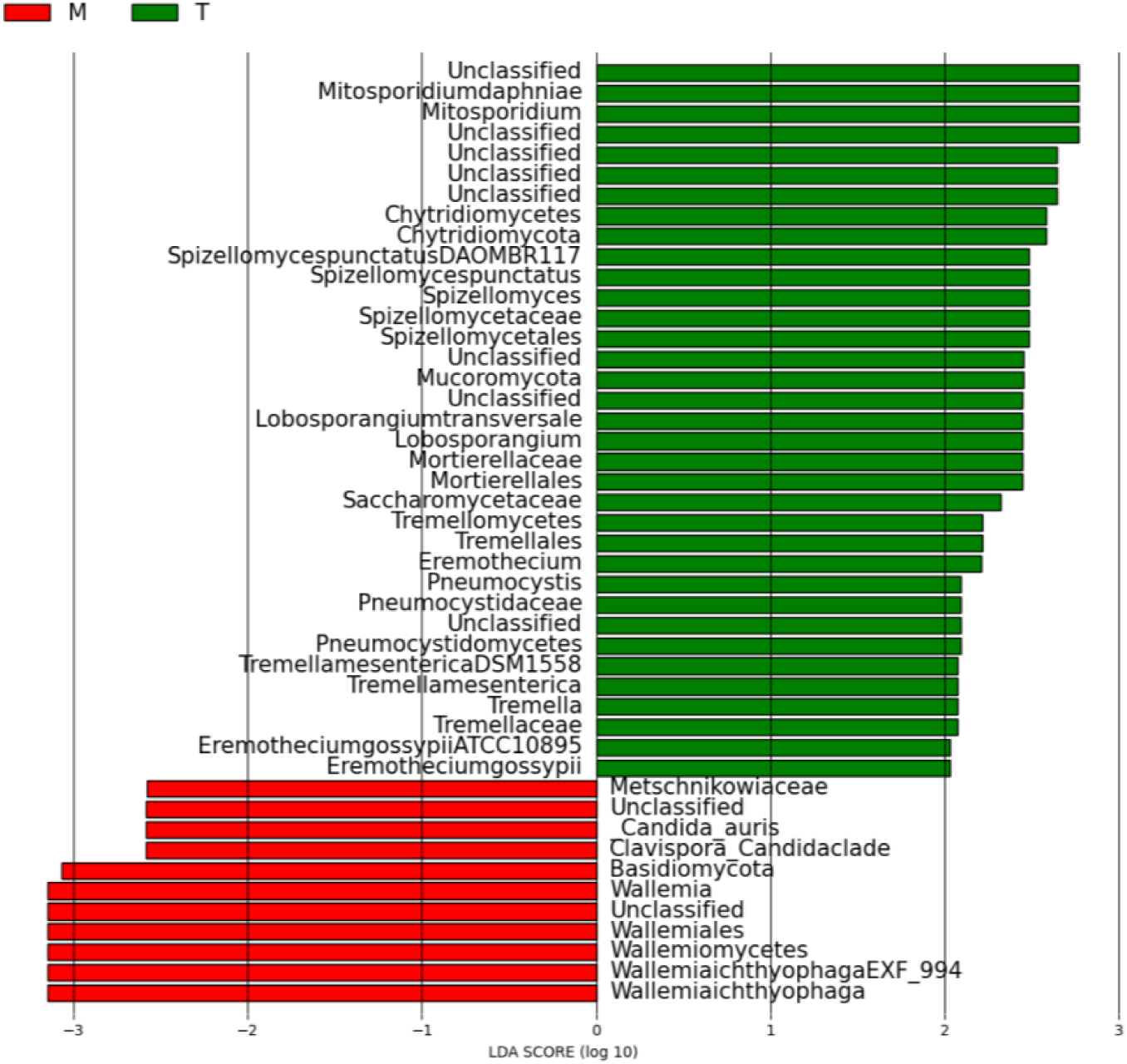
Clades identified to be explanatory of the difference between fungal community profiles in *D. majalis* and *D. traunsteineri* environments using linear discriminant analysis effect size (LEFSe). Only clades with a minimum log(LDA) score of 3 and a Kruskal-Wallis p-value < 0.05 are shown.

**Supplementary Figure 3.**
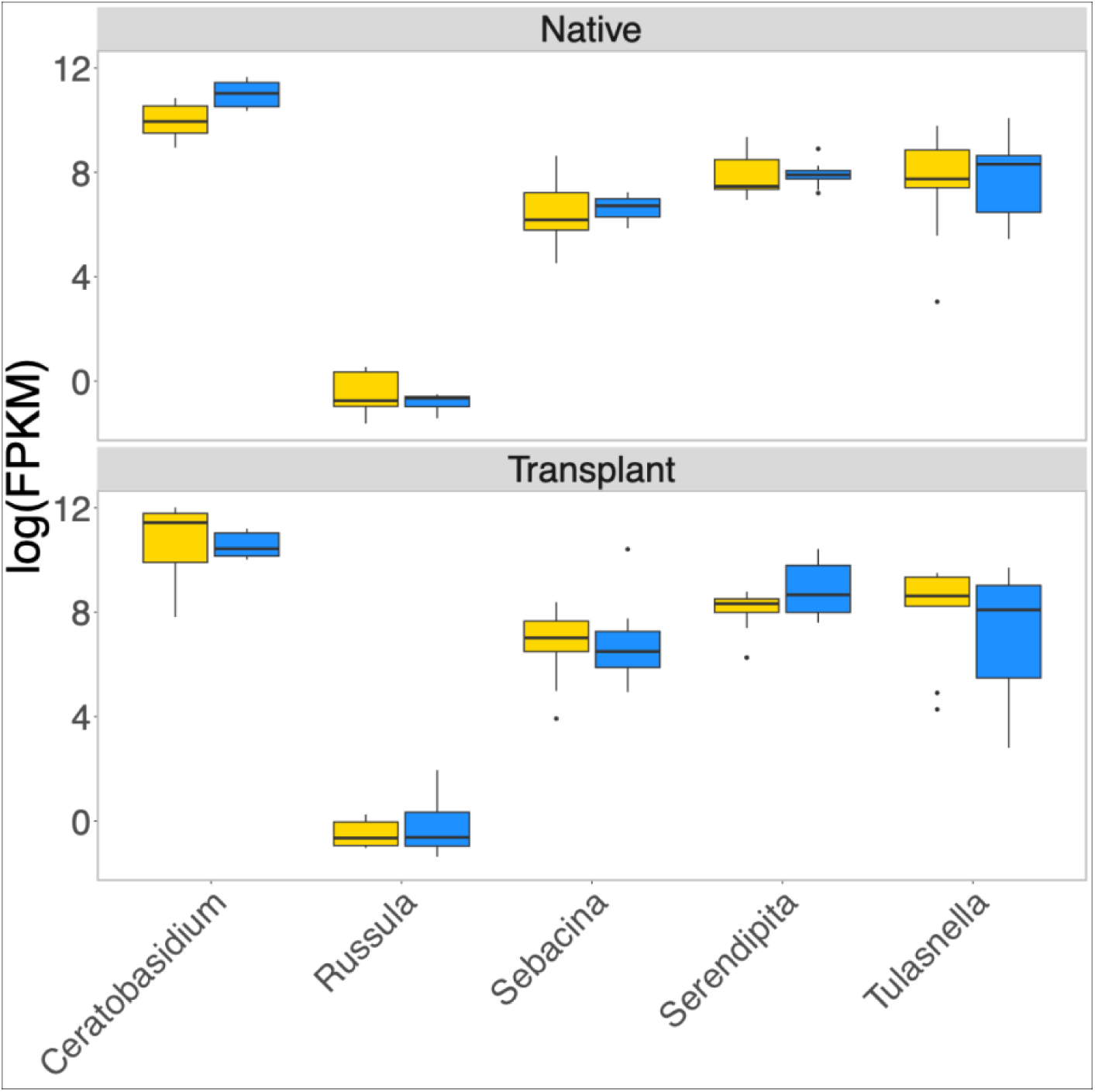
Boxplot showing FPKM normalised read counts mapping to representative ITS sequences of five selected fungal taxa reported to form mycorrhizal associations with orchid species. FPKM normalised read counts are shown on a log scale.

**Supplementary Figure 4.**
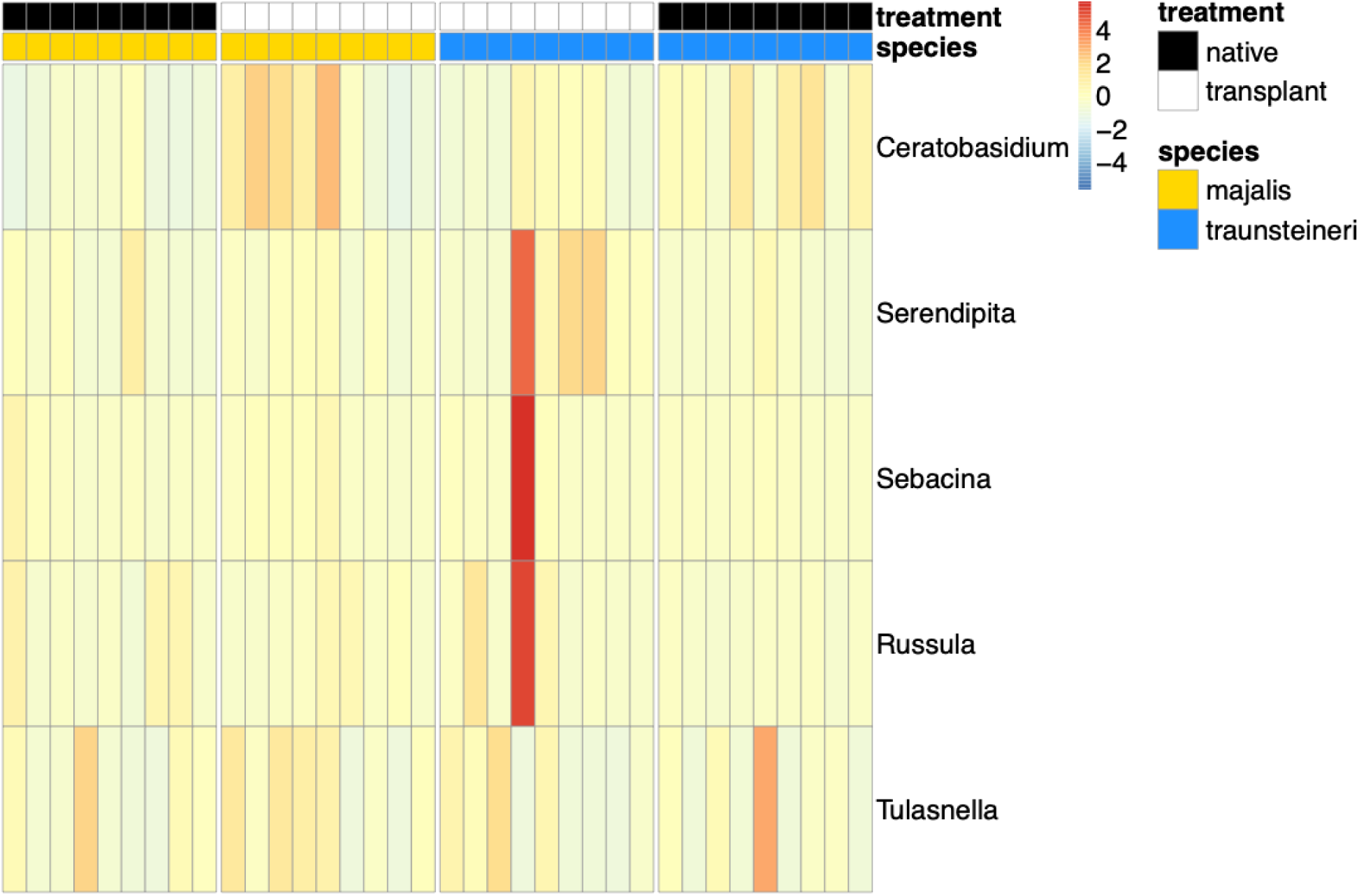
Heatmap of FPKM normalised ITS read counts across all samples of *Dactylorhiza majalis* and *D. traunsteineri*. Treatment of samples (native or transplant) is shown in black and white on the top annotation bar, and species is shown in blue and yellow on the bottom annotation bar. FPKM normalised counts are scaled prior to visualisation.

