## Supplementary Figure 1 for "Ecological divergence of sibling allopolyploid marsh orchids is associated with species specific plasticity and distinct fungal communities"

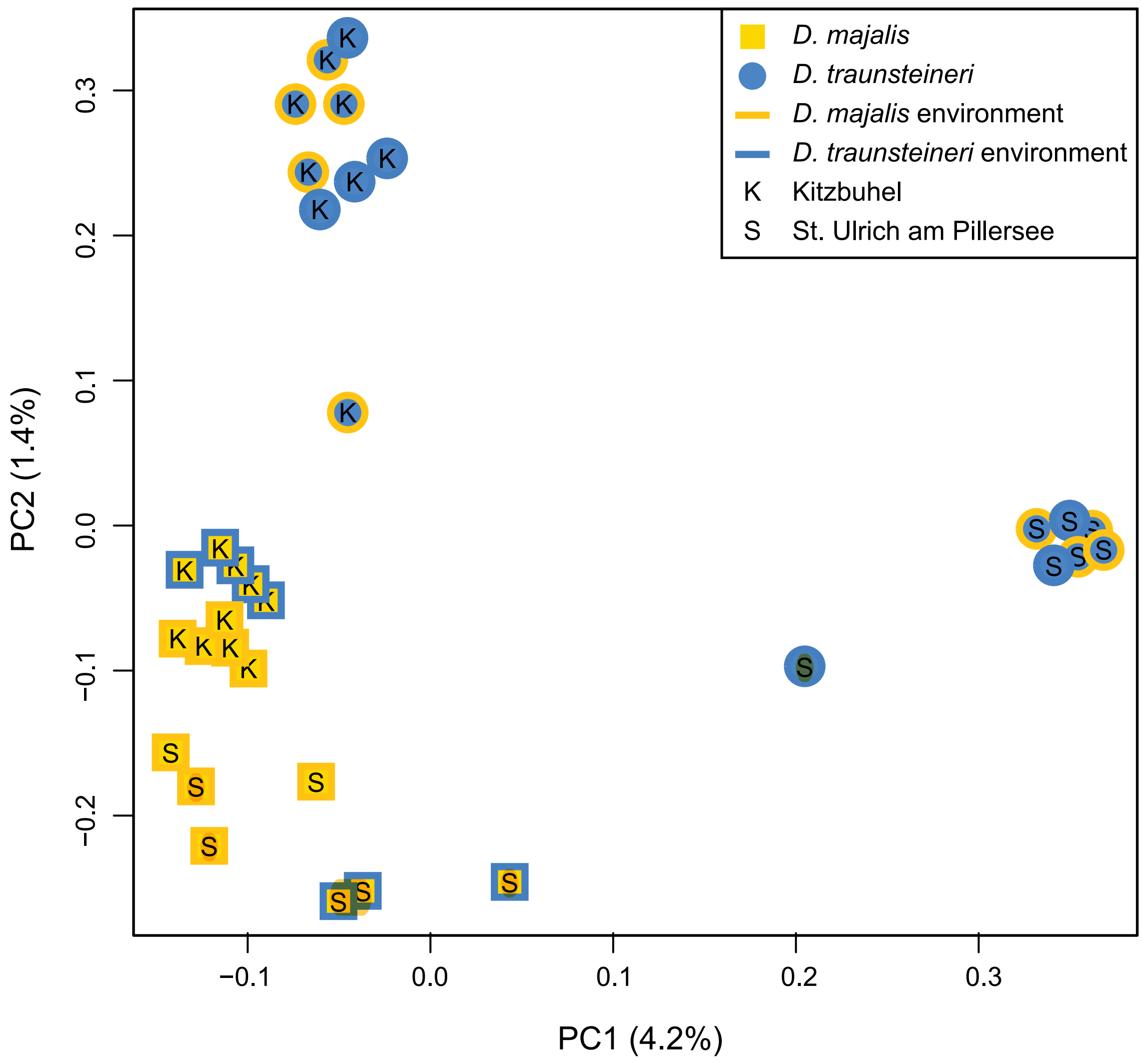


Supplementary figure 1. PCA plot showing genotypic variability in Dactylorhiza majalis and D. traunsteineri in the two localities tested (Kitzbuhl and St. Ulrich). Points represent replicates of D. majalis (yellow filled symbols) in native and D. traunsteineri (blue filled symbols) in native and transplanted habitats (according to the legend).
