## Supplementary figures and images for "Ecological divergence of sibling allopolyploid marsh orchids is associated with species specific plasticity and distinct fungal communities"

### Supplementary Figure 2

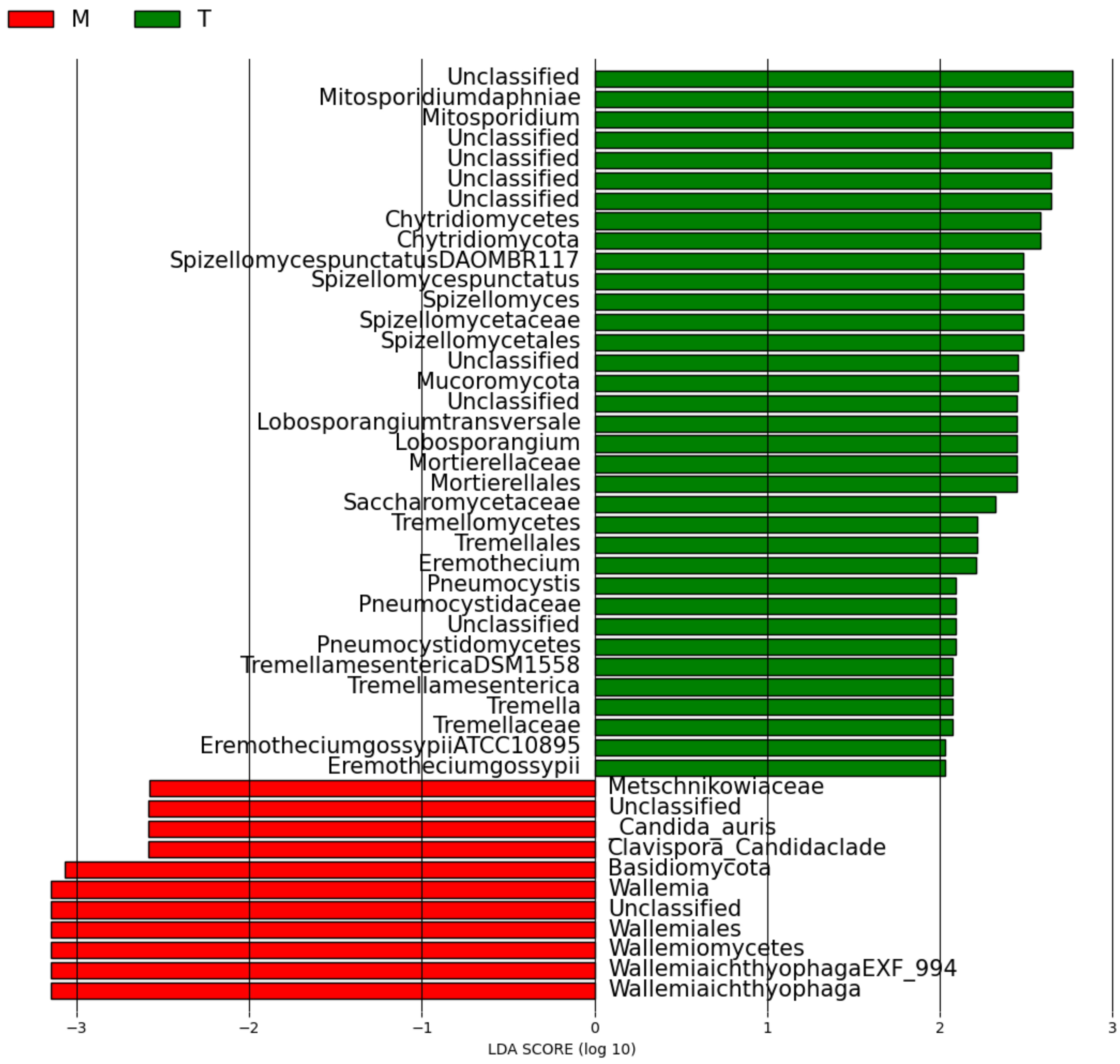
